# Prospective Motion correction improves the sensitivity of fMRI pattern decoding

**DOI:** 10.1101/296442

**Authors:** Pei Huang, Johan D. Carlin, Arjen Alink, Nikolaus Kriegeskorte, Richard N. Henson, Marta M. Correia

## Abstract

We evaluated the effectiveness of prospective motion correction (PMC) on a simple visual task when no deliberate subject motion was present. The PMC system utilizes an in-bore optical camera to track an external marker attached to the participant via a custom-moulded mouthpiece. The study was conducted at two resolutions (1.5mm vs 3mm) and under three conditions (PMC On and Mouthpiece On vs PMC Off and Mouthpiece On vs PMC Off and Mouthpiece Off). Multiple data analysis methods were conducted, including univariate and multivariate approaches, and we demonstrated that the benefit of PMC is most apparent for multi-voxel pattern decoding at higher resolutions. Additional testing on two participants showed that our inexpensive, commercially available mouthpiece solution produced comparable results to a dentist-moulded mouthpiece. Our results showed that PMC is increasingly important at higher resolutions for analyses that require accurate voxel registration across time.

## INTRODUCTION

One of the biggest weaknesses in functional magnetic resonance imaging (fMRI) is arguably the long scanning times, typically in the range of 5 to 15 mins per scan to ensure sufficient statistical power (Birn et al., 2013; Murphy et al., 2007). This results in a wide array of problems, most notably the degradation of data quality due to subject motion. As the signal of interest in most fMRI studies is small (Renvall et al., 2014; Runeson et al., 2013), any decrease in data quality could obscure the signal. In addition, motion correlated to the stimulus has also been shown to result in false-positive activations (Field et al., 2000; Hajnal et al., 1994).

Methods to correct for motion can be broadly classified into Retrospective Motion Correction (RMC) and Prospective Motion Correction (PMC). Historically, the use of RMC has been more widespread due to convenience and limited ability to acquire time-linked motion data in the MRI scanner. In RMC, rigid body translations and rotations are applied to each volume post-scan to align all acquired volumes to the same scan (Ashburner and Friston, 2003; Johnstone et al., 2006). Although this works well for slow motion between acquisitions, RMC is unable to correct for spin history effects and k-space distortion due to intra-volume motion (Goebel et al., 2006; Penny et al., 2011). Moreover, as the acquisition box is not coupled to the brain, edge voxels could be lost due to subject motion in the case of partial brain acquisitions. These drawbacks become increasingly important at higher resolutions and higher fields because the field of view is typically more restricted.

The field has been shifting towards PMC as a way to reduce the impact of subject motion. PMC requires an acquisition of the movement parameters of the participant’s head concurrently with the acquisition of the imaging volume (Callaghan et al., 2015; Maclaren et al., 2012). The motion parameters are used to update the position of the acquisition box within the participant’s head just before each radio-frequency (RF) pulse. Maclaren et al., 2013 and Zaitsev et al., 2016 provide a good overview on the current state of the field and list the most promising techniques, some of which have demonstrated significant benefits to data quality relative to RMC (Muraskin et al., 2013; Stucht et al., 2015; Todd et al., 2015). The estimation of PMC parameters can be done by either using the internal MR data or external tracking modules. Internal MR data methods, such as k-space navigators (Van Der Kouwe et al., 2006; Ward et al., 2000) or fat-based navigators (Engstrom et al., 2015), require additional scans between each acquisition, which would reduce the temporal resolution of the data further. External tracking modules, including the system we evaluate here, utilize a secondary system to acquire the positional data in real time and transfer the data to the scanner.

For 2D Echo-Planar Imaging (EPI) sequences, we expect PMC to improve the quality of the data relative to RMC. First and foremost, PMC allows for slice-wise realignment instead of whole volume realignment. This allows for correction of both intra and inter-volume motion, while conventional RMC implementations only correct for inter-volume motion. Secondly, accurate coupling of the acquisition box to the participant’s head removes spin history effects and preserves edge voxels (Yancey et al., 2011). This allows for a higher confidence in voxel-wise registration across volumes and ensures that the entire acquisition volume can be used for model fitting.

One of the major problems when implementing PMC using optical tracking is the method of attachment of the marker to the participant. There is a general consensus that skin attachment is insufficiently rigid (Callaghan et al., 2015; Muraskin et al., 2013; Stucht et al., 2015; Todd et al., 2015) and similar conclusions were drawn from our own initial testing (Huang et al., 2017). Some sites use dentist-moulded mouthpieces to ensure perfectly rigid coupling (Stucht et al., 2015). However, this is both time-consuming and expensive as participants are required to visit a dentist at least a day prior to the actual scan to provide a mould which is used to make the custom mouthpiece. Here, we attempt a novel method of moulding the mouthpiece on the spot. This reduces both time and monetary costs, which could in turn allow PMC to be adopted more widely. As part of our study, a comparison between our novel method and a dentist moulded mouthpiece was also carried out on two participants.

Most previous studies that investigated the effectiveness of PMC used deliberate subject motion (Ooi et al., 2011; Schulz et al., 2014; Todd et al., 2015), which may not be representative of actual participant behaviour in the scanner. For instance, deliberate motion is likely to result in larger head displacements than what would be observed in a typical participant instructed to remain still, and thus result in an overestimation of the benefit of PMC. Moreover, most studies have been focused on structural scans, with fewer studies examining functional scans. Todd et al., 2015 utilized an optical tracking system to correct 3D EPI resting state data under three separate conditions, no motion, slow deliberate motion and fast deliberate motion. They demonstrated that PMC application significantly increased tSNR of resting state fMRI in both motion conditions, while appearing not to affect the data for scans with no motion. In the same paper, they also demonstrated an increase in significant voxels for a motor and visual task, albeit only in a single subject. Another group utilized the same system for 2D EPI and did not observe any improvement in tSNR values for a finger tapping task (Zaitsev et al., 2016). The authors noted that this is likely due to the poor adhesion of the marker as the marker was attached to the nose instead of via a mouthpiece. For task-based fMRI, Rotenberg et al., 2013 demonstrated that using a stereo optical tracking setup can significantly reduce false-positive rates in actively moving participants. Schulz et al., 2014 adopted an experimental design where participants were asked to keep their head still while moving their legs and also demonstrated a reduction in false positives. While no deliberate head motion is carried out by the participant, task correlated motion is still present and hence, unlikely to be representative of typical participant behaviour in the scanner.

In the present study, we evaluated the effectiveness of PMC with a custom mouthpiece attachment on a typical fMRI experiment, where participants were asked to remain as still as possible for the duration of the scan. We focused on the benefits of PMC in preserving information in fMRI response patterns. Visual gratings were chosen as stimuli because their encoding in visual cortical response patterns is reasonably well understood. In particular, we examined the accuracy of multivariate decoding in the primary visual cortex (V1), which is generally high (Alink et al., 2013; Kamitani and Tong, 2005; Tong et al., 2010).

A further advantage of our experimental stimuli is that they may enable analysis of the effect of PMC on multiple spatial scales. Human V1 representation of visual stimuli occurs over different spatial scales: There is a general, coarse selectivity pattern due to radial bias, and additional selectivity on a finer spatial scale that is independent of radial bias. Such fMRI effects may originate in the topography of underlying neuronal population codes. Specifically, neurons responding to radial orientations with respect to the fixation point appear to be more frequent, creating a global areal map of radial orientation frequencies (Freeman et al., 2013; Sasaki et al., 2006). Neurons responding independently of radial bias are organised in a much finer-grained columnar map of orientation preference (Alink et al., 2013; Swindale et al., 2003; Tong et al., 2010). The different spatial frequencies of these two nested organisations leads us to expect that that the effectiveness of PMC may vary with the spatial scale of the fMRI measurement, and with the visual field coverage of a given region of interest. Specifically, we expect a maximal benefit of PMC in higher-resolution acquisitions and for regions of interest that do not include expected radial bias signal.

For this study, Linear Discriminant Contrast (LDC) was chosen as the primary metric for investigating multivariate effects. The LDC is a continuous statistic derived from the well-known Fisher’s linear discriminant. As in conventional linear discriminant analysis, it uses the training data to generate a set of representative weights for each voxel to maximize sensitivity between the two conditions. These weights are applied to the testing data to form the LDC, which serves as a measure of the reliability of the difference between the two conditions across the training and testing data. For completeness, results from a more conventional Support Vector Machines (SVM) classifier were also included. In SVM, the training data is used to determine the decision boundary between the two categories. The testing data is then mapped onto the same space and assigned a category based on their position with respect to the boundary.

Both methods use a linear decision boundary that is described by a weights vector, although the generation of these weights differs between methods and LDC is expected to be more sensitive due to the following three differences between the two methods (Walther et al., 2016). Firstly, the SVM classifier discretizes the output into a binary system of either 1 for correct or 0 for incorrect. This reduces the sensitivity of the analysis to small fluctuations in the data. Secondly, the SVM classifier suffers from ceiling effects where the performance is capped at 100%. Lastly, in LDC, we take a dot-product between the weights vector and the contrast estimate from the test data. This generates an estimate of the coherence of the data without needing to establish a threshold parameter. In contrast, an absolute threshold is used in SVM to classify the data into the two groups. As the data in fMRI are unitless and the magnitude of responses can shift significantly between runs, it is plausible that classification errors could result from an SVM classifier that had learned the correct weights for the voxels, but applied an incorrect threshold due to between-run variations of response magnitudes.

## METHODS

### Prospective Motion Correction (PMC)

The PMC system employed here utilizes an in-bore optical camera (Kineticor, HI, USA) to track the motion of a passive Moiré phase marker at a frame rate of 80Hz (Maclaren et al., 2012; Weinhandl et al., 2010). Layered gratings and the design of the marker allow all three translational and three rotational degrees of freedom to be measured. The precision of the translational and rotational measurements were previously reported to be 0.1mm and 0.1 degrees respectively (Maclaren et al., 2012). This information is logged to the PMC system and transferred to the scanner host computer. This is then transformed from camera to scanner co-ordinates using a calibrated transformation matrix acquired pre-scan. The parameters in scanner co-ordinates are then used to update the imaging gradients, RF frequency and phase for each repetition time (TR) to ensure that the same imaging field-of-view (FOV) is acquired across TRs (Herbst et al., 2012; Speck et al., 2006). This update process requires an accurate marker-to-brain motion coupling. To that end, a custom mouthpiece (Figure 1) was made for each participant before each session. Dental putty (Provil Novo: Putty Fast) was mixed and loaded onto a dental impression tray (Tra-Tens^®^ Impression Trays, Waterpik). Participants were asked to bite on the tray for 2 minutes to allow the putty to harden. Once moulded, the tray remains firmly attached without requiring active biting from the participant. The marker was attached to the tray via a 3D printed plastic arm with 3 pivot points to allow flexible positioning of the marker within the field of view of the tracking camera.

**FIGURE 1:**
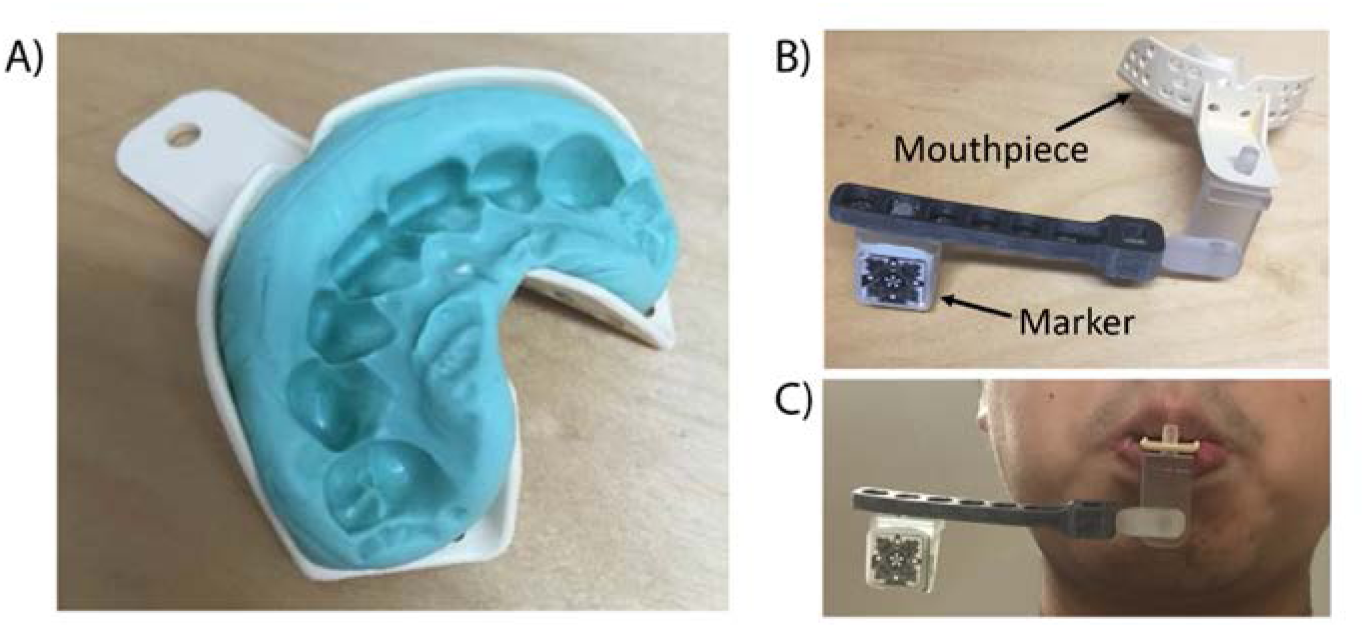
A) An example of moulded and hardened dental putty in the shape of the participant’s teeth. Dental putty (Provil Novo: Putty Fast) was used and once hardened, no deliberate effort was required from the participant to keep the mouthpiece in place. B) The marker is attached to the mouthpiece via an arm extension with 3 pivot points to allow for flexible positioning of the marker. C) A sample image of the entire setup when attached to a participant.

### Experimental Design

We adopted a 2x3 factorial design for data acquisition: 1.5mm isotropic voxels vs 3.0mm isotropic voxels and PMC On, Mouthpiece On (P+M+) vs PMC Off, Mouthpiece On (P-M+) vs PMC Off, Mouthpiece Off (P-M-). The fourth permutation, PMC On, Mouthpiece Off cannot be tested as the mouthpiece was required to acquire the positional information for PMC. In condition P-M+, while PMC was not applied to the MRI data, tracking data were still acquired.

The three separate scan conditions allowed for isolation of the following experimental effects: Comparing data from conditions P+M+ and P-M+ demonstrates the impact of PMC correction, while controlling for the presence of the mouthpiece, and comparing data from conditions P-M+ and P-M-quantifies the effect of the mouthpiece. Most importantly, comparing the data from conditions P+M+ and P-M-showcases the net benefit of implementing PMC in actual studies.

Data analysis was carried out over 3 distinct ROIs, the entire V1, regions with radial bias and regions without radial bias. Regions with radial bias are expected to have more coarse-grained response patterns and hence, should be more robust against motion effects. In contrast, regions without radial bias should have more fine-grained response patterns and be more sensitive to motion effects.

### Data Acquisition

All scanning was performed on a Siemens 3T Prisma-Fit scanner using a standard 32-channel head coil. Participants provided informed consent under a procedure approved by the institution’s local ethics committee (Cambridge Psychology Research Ethics Committee). A total of 18 healthy participants were scanned (8 females, age range 20-41, 1 participant was an author of this study).

Each participant was present for three repeat sessions under each of the three conditions, P+M+, P-M+ and P-M-. All other scan procedures and sequences were preserved across sessions, but the order of the conditions was randomised across participants. In all cases, participants were instructed to remain as still as possible so as to mimic a typical fMRI experiment. Participants were blinded as to whether PMC was applied (P+M+ vs P-M+) to prevent bias but were aware when no mouthpiece was present (P-M-). The interval between sessions was not controlled due to restrictions imposed by participant and scanner availability. The range of intervals between sessions was 1 to 20 days.

For each session, MPRAGE structural images were acquired first (TR=2250ms, TE=2.22ms, TI=900ms, GRAPPA=2, FOV=256mm*256mm*192mm, Matrix size=256*256*192, FA=9°, ToA=˜5mins). This was followed by a total of four functional task scans: two main experimental scans and two localizer scans. For the main experimental scans, the participants were scanned while viewing the gratings in a block design (Figure 2), once each at voxel resolutions of 3mm and 1.5mm. The data from these scans were used to compare the data quality across conditions. The acquisition order for the two resolutions was randomised across participants, but remained constant for the three sessions for the same participant. The two localizer scans were carried out at 3mm resolution and used to generate a retinotopic map for the segmentation of Regions of Interest (ROI). Each session was followed by an eight minute resting state scan for each participant. Upon completion of scanning for each session, the participants were asked to fill out a short questionnaire with regards to the comfort of the mouthpiece.

**FIGURE 2:**
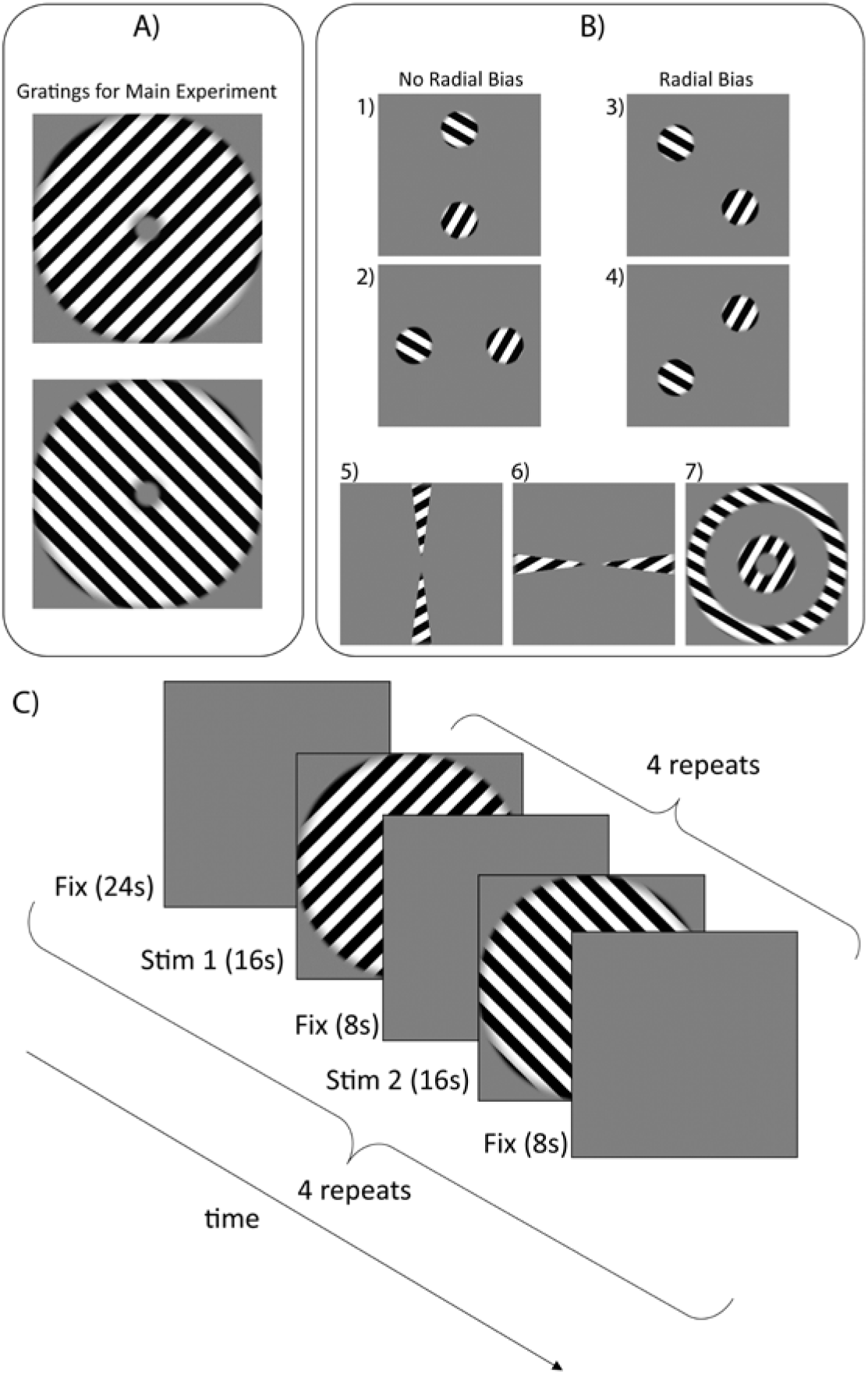
A) Uniform gratings of two different orientations (both 45° from the vertical) were used as stimuli for the main experiment. B) The seven stimuli were presented in a randomized block order during localizer scans. Stimuli 1 and 2 were used to isolate regions with no radial bias, Stimuli 3 and 4 were used to isolate regions with radial bias and stimuli 5, 6 and 7 were used to segment V1. C) An illustration of the timecourse of stimuli presentation for the main experiment. The two 16s stimulus blocks (one for each orientation) were repeated four times each for a total of eight stimulus blocks per sub-run. The entire sub-run was repeated four times for each scan, with a gap of 24s fixation between sub runs to minimise the dependency between sub-runs.

Field-of-view (FOV) parameters for both 3mm and 1.5mm EPI sequences were chosen such that the same volume (192mm*192mm*90mm) was imaged across scans. Imaging parameters for the 3mm isotropic EPI were: TR=1260ms, TE=30ms, FA=78°, Matrix size=64*64*20, ToA=˜11mins. Imaging parameters of the 1.5mm isotropic EPI were: TR=3050ms, TE=30ms, GRAPPA=2, FA=78°, Matrix size=128*128*40, ToA=˜11mins. Imaging parameters for the 3mm resting state EPI were: TR=2000ms, TE=30ms, FA=78°, Matrix size=64*64*32, ToA=˜8mins.

### Stimulus Design

All stimuli were created using Matlab (2009a, The MathWorks, Natwick, MA, USA) and presented in the scanner using Presentation (v17.2). All stimulus types were presented within an annulus (inner radius = 1.05°, outer radius = 7.15°) centred on fixation on a mid-gray background. For the main experiment, uniform grating stimuli (45° clockwise or 45° anti-clockwise from the vertical) were used (see Figure 2, panel A) as they are balanced about both vertical and horizontal orientations. Thus a global preference map for these orientations will yield an equal global activation pattern for each grating (Furmanski and Engel, 2006; Seymour et al., 2010). The gratings had a spatial frequency of 1.25 cycles per visual degree to strongly drive responses in V1 (Henriksson et al., 2008). For each orientation, 20 different stimuli were generated with spatial phases uniformly distributed between 0 to 2π.

Both orientations of the gratings were presented in a block design within each run. Each run was divided into four equal sub runs, which contained eight 16s stimulus blocks each. Stimuli from each orientation were presented in an alternating order, with an alternate leading orientation across sub runs (see Figure 2, panel C). Within each block, the 20 phase-shifted stimuli of one orientation were presented in a randomized order at a frequency of 2 Hz. The stimulus duration was 250ms, followed by 250ms of fixation. Each block was separated by a 8s fixation period and each sub run was separated by a 24s fixation period. This ensured that estimates obtained from each sub run are independent from each other.

In order to define regions of interest (ROIs) for all of V1 and V1 sub-regions with and without radial bias, localizer scans were conducted for retinotopic mapping. We presented dynamic grating stimuli designed to optimally drive responses in selective regions of retinotopic early visual cortex. Seven such stimuli groups were used (see Figure 2, panel B): (1, 2) a patch pair stimulus consisting of two circular patches (spanning 2.40°–5.80° eccentricity) lying along the vertical or the horizontal axis, respectively; (3, 4) a patch pair stimulus of the same kind lying along the two diagonals, respectively; (5) a horizontal double-wedge stimulus, spanning a polar-angle range of ± 15° around the horizontal meridian; (6) a vertical double-wedge stimulus of the same kind; (7) a 1.5°-wide ring peripherally surrounding the main-experimental stimulus annulus (5.65°–7.15° eccentricity), and a 1.5°-wide ring inside the annulus (1.05°–2.55° eccentricity). Each stimulus group contained rectangular gratings with the same spatial frequency as the main gratings. For each stimulus group, the orientations of the rectangular-phase gratings were randomized in angular steps of π/6, and for each orientation, there were four different spatial phases for the stimuli. The stimuli were presented in 13s blocks. The four patch pair stimuli were presented eight times each while the other stimuli were presented four times each in a randomized sequence over a run lasting 8 minutes. This was repeated twice per session for each participant.

During all runs, including the functional localizer scans, participants were instructed to continuously fixate a central blue dot (diameter: 0.1° visual angle). At randomized time points during the experiment, the dot flashed green for 250ms at an average rate of once per 3.5s (with a minimum gap of 1.5s between colour changes). The participants were tasked to press a button with their right index finger in response to every flash to encourage fixation. Task accuracy was calculated by dividing the number of flashes that the participant responded to within 2 seconds by the total number of flashes. Three participants with lower than 50% response accuracy on the task for any run were completely excluded from further analysis.

### Data analysis

The goal of this study was to evaluate the performance of the PMC system for a standard fMRI experiment and its impact on the quality of the data obtained. In order to accomplish this, several different metrics were employed to study the imaging data quality.

The first three image volumes for each scan were discarded to allow for the signal to reach steady-state. The data sets were processed in a standard pipeline using SPM8 (Penny et al., 2011), which included rigid-body realignment to correct for head motion followed by temporal sinc interpolation to correct for differences in slice acquisition times. Linear and first order sinusoidal detrending were applied to the data to remove signal drift.

### Analysis of SPM motion parameters

The realignment parameters were extracted from SPM and collated. In condition P+M+, this measure indicates the amount of residual motion that PMC failed to correct. Comparing condition P-M+ and condition P-M-allowed for quantification of the impact of the mouthpiece. To combine data from all six degrees of freedom into one integrated motion metric per scan, rotation angles were converted into displacement measures using a rotational radius of 5.7cm (which is reasonable considering the typical head size of an adult, Todd et al., 2015), and the square root of the sum of squares of the resulting six displacement parameters were calculated per unit time.

### Regions of Interest (ROI)

All ROI segmentations were done in Freesurfer 5.3.0. Activation t-maps were obtained in SPM by fitting a GLM to the fMRI data from the localizer run. The maps were projected onto polygon-mesh reconstructions of individual participants’ cortices. For this study, V1 served as the main region of interest. The boundaries of V1 were obtained by contrasting the t-maps for the vertical wedges against horizontal wedges and contrasting the t-maps for the localizer rings against all four patch pairs.

Further analysis was done by segmenting regions activated by patch pairs 1 and 2 and patch pairs 3 and 4 (Figure 2, Panel B). Boundaries for each patch pair were obtained by contrasting the patch-pair of interest against all other patch pairs and the localizer rings (see Supplementary Figure S1). As both orientations of the grating stimuli are at an angle of 45 degrees with respect to the axis joining the centre of the patch to the centre of the stimuli for patch pairs 1 and 2, there should be minimal effect of radial bias for these regions. In contrast for patch pairs 3 and 4, the grating stimuli lie either perpendicular or parallel to the axis on which the patch pair lies, hence resulting in maximal radial bias. Due to the difference in spatial frequency of the activation patterns, regions driven by radial bias are expected to be more robust against motion effects. When the whole V1 is employed for classification training, regions responding to radial bias would be expected to strongly drive classification performance. This could mask subtle differences in the data arising from small amounts of motion. Hence, data analysis was carried out on regions with and without radial bias individually, as well as the entire V1.

### Temporal Signal to Noise Ratio (tSNR) analysis of rsfMRI

A whole brain mask was generated using the brain extraction tool (Smith, 2002) in FSL (Jenkinson et al., 2012). The rsfMRI underwent slice-time correction and realignment prior to application of the brain mask to extract all voxels within the brain. For each voxel, the tSNR was obtained by dividing the mean voxel intensity across the entire resting-state time course by the standard deviation of the voxel intensity.

### Univariate analysis using fCNR

Functional Contrast-to-Noise Ratio (fCNR) was calculated for the entire V1 from the main experiment. The post-processed fMRI data were fitted to a simple GLM which only modelled whether a stimuli was present or only the fixation dot was present. The fCNR was obtained by dividing the amplitude of the signal by the standard deviation of the noise. The amplitude, in this case, is obtained by taking the absolute difference between the baseline of the signal and the signal peak (Welvaert and Rosseel, 2013). The signal peak was obtained by multiplying the contrast estimate by the GLM and taking the maximum value. Note that zero corresponds to the baseline, since non-stimulus periods were not modelled (ie baseline was modelled implicitly in the GLM).The standard deviation of the noise was obtained by taking the standard deviation of the residuals of the GLM.

### Cross-validated Linear Discriminant Contrast (LDC) Analysis

In this study, the cross-validated LDC (Kriegeskorte et al., 2007; Walther et al., 2016) between the responses to the two different orientations is used as the metric for investigating multivariate effects. Similar to a univariate test, the LDC is effectively a contrast between two conditions measured on a discriminant. This discriminant is made up of a weighted combination of the ROI voxels, where the weights have been chosen with independent data to produce maximum sensitivity to the difference between the two conditions. Cross-validating the contrast removes the positive bias affecting estimates of distances (which are by definition positive) from noisy data (Walther et al., 2016). This measure is also known as the crossvalidated Mahalanobis (crossnobis) distance (Kriegeskorte and Diedrichsen, 2016).

For the purposes of LDC analysis, all presentations of each orientation, after the extraction of one-sub-run, were modelled as one event in the design matrix. This forms the independent dataset used to generate the LDC weights. In the extracted sub-run, all presentations of each orientation were also modelled as one event. Because LDC is a continuous statistic, extraction of separate response pattern estimates for each block was not needed, unlike classification accuracy. This provides an additional benefit over SVM as modelling using a single event for all presentations provides a more stable estimate of the activation pattern (Abdulrahman and Henson, 2016). We also showed that repeating the LDC analysis with individual blocks modelled produces less sensitive results (see Supplementary Figure S2, Supplementary Table S3).

For each participant, data from three sub-runs were used to generate a representational distance metric by fitting the detrended data to the detrended design matrix and calculating a pairwise contrast between the responses to the two orientations. This representational distance metric was normalized using the sparse covariance matrix (Ledoit and Wolf, 2003) of the noise residuals to produce a weights vector. Next, we calculated the LDC test statistic by taking the dot product between the weights vector and the contrast estimate from the test sub run. We repeated this procedure for each of the four possible sub run cross-validation iterations, and averaged over the LDC statistics to obtain a final continuous performance estimate, which is centred on zero under the null hypothesis of no reliable response pattern difference between the stimuli. As the number of voxels used in this analysis varied across regions, resolutions and conditions, the LDC was normalized by dividing the metric by the square root of the number of voxels.

### Support Vector Machines (SVM) classifier

SVM classifiers are a type of supervised machine learning algorithm that can be trained on a labelled sub-set of the data to classify the remaining unlabelled data. The predictions from the SVM classifier are compared to the actual labels to calculate the accuracy of the classifier. All SVM classification was done using the SVM classifier in the Matlab Bioinformatics toolbox. While we have presented reasons on why LDC is the better analysis method for analyses that depend on detecting differences in discriminability between sets of conditions, we included the results from the SVM classifier as it is currently the more commonly used method.

For this study, each 16s block of stimuli was modelled as an individual epoch to generate a design matrix comprising 16 regressors per stimulus orientation. This modelling setup was used to provide a larger number of samples for the training and testing data sets, which would allow for more stable estimates of classification accuracy. The processed fMRI data were fitted to the design matrix using ordinary least squares to obtain beta values (fitted parameter estimates for each voxel and presentation block).

The beta value maps were split into two groups, labelled by the orientation of the stimuli. All presentations of the stimuli from three sub runs were used to train the classifier. The classifier was then tested on the eight presentations that were not used for training. This was reiterated four times per participant per condition per resolution, leaving a set of presentations out of the training each time (leave-one-sub-run-out cross-validation). We take above-chance accuracy over the splits to indicate a persistent representation of the stimuli across time, and changes in classification accuracy to indicate effects of PMC on the data quality.

### Simulations for Comparison of MVPA Methods

To allow for a more informed comparison between the two MVPA methods’ sensitivity to PMC-like effects, a set of computational simulations was also conducted and included in Supplementary Figure S4. This simulation was designed to closely resemble real dataset’s design (alternating block design, 16s per block, 8 blocks per sub run, 4 sub runs per participant, 15 participants in each iteration), and analysis approach.

We first generated a design matrix with two regressors (one regressor per stimulus orientation). For each of 100 iterations, we multiplied this design matrix by a contrast vector to simulate the activation timecourse of a voxel. The activation response to each stimulus orientation was obtained by sampling from independent Gaussian distributions of with mean 0 and standard deviation 1. This generates a contrast vector centred around 0, with an expected mean absolute difference between conditions of 2/sqrt(pi). This expected mean is calculated by integrating abs(x1-x2)P(x1)P(x2) across all possible values of x1 and x2. This was normalized to give a mean absolute contrast of 1 and repeated 500 times (average number of voxels at 3mm) to create a simulated, noiseless fMRI dataset for 1 participant. We repeated this procedure separately for each of 15 simulated participants.

Three types of noise were added to each simulated participant’s data. Firstly, at the level of the activations, independent Gaussian noise (mean 0, standard deviation 1) was added to each presentation block to reflect variations in attention to the stimuli by the participant for each voxel. Secondly, thermal noise was modelled using independent Gaussian noise for each voxel and each timepoint (mean 0, standard deviation 1.8). The first two sources of noise were assumed to be constant in variance across all conditions. Lastly, the physiological noise, arising due to heartrate, respiration and head motion, was modelled by generating 10 independent Gaussian noise vectors for the entire timecourse (mean 0, standard deviation 1), and projecting a randomly weighted combination of these 10 vectors onto each voxel. The underlying physiological noise time courses were independent across sub runs, but the projection onto voxels was held constant for a given participant, thus providing a reliable covariance structure that the discriminant methods could exploit. We varied the physiological noise level to simulate the effect of PMC on our data. This simulation approach assumes that the effect of PMC is to reduce physiological noise in the data, without affecting other noise sources.

To estimate the amount of thermal noise in our model, we first estimated the thermal noise in our real data by measuring the fluctuations in signal intensity outside the brain. The ratio of thermal noise to total noise in our real data was then calculated and used to generate as estimate for the thermal noise in our simulations. Using fCNR of 1.8 (similar to that of the 3mm data), we then estimate the thermal noise in our simulations to be a standard deviation of 1.8.

Each simulated participant’s data were then passed through both LDC and SVM analysis. Simulated group-level differences in LDC or discriminant performance were then assessed with pairwise t-tests were conducted across varying fCNR. Finally, we calculated rejection probability for each fCNR pairing as the proportion over the 100 simulated datasets where statistical significance was attained (p<0.05, Supplementary Figure S4). The thermal and physiological noise sources were scaled to achieve two objectives: First, a mean fCNR over voxels that matched the real data (namely a fCNR of 1.6 for 1.5mm data and 1.8 for 3mm data); Second, a range of fCNR differences (ie, strength of physiological noise manipulation) that made it possible to observe the full range of rejection probabilities. Note that these values of fCNR are distinct from the fCNR calculated in the previous section, because we are now defining the contrast as the difference in activation between the two stimulus orientations, rather than presence vs absence of stimuli as in the previous section. Both simulation and real data also had a 100% overall LDC vs chance rejection probability.

### Permutation Testing for Significance

As most of our data (motion parameters, tSNR and fCNR) does not satisfy the continuous or normality assumptions of standard parametric tests, pairwise permutation tests were carried out on pairs of these data to test for significance. For each iteration of a pair of conditions, the labels for the measures used were randomized within each participant and the mean difference between conditions recorded. Each pairing was iterated 10000 times to generate a distribution and the actual mean difference obtained from the study was tested against this distribution. This permutation test models participant as a fixed effect (FFX), and produces similar p values as a fixed-effect T test when Gaussian assumptions hold.

### Three-way repeated measures ANOVA

The LDC data was analysed using a three-way repeated measures ANOVA, given that the data can be assumed to be continuous and approximately normally-distributed. Post-hoc comparisons were done using the Tukey’s honest significant difference (HSD) test, which corrects for multiple comparisons. The repeated measures ANOVA models participants as a random effect (RFX), and can thus support inferences about the sampled population.

### Comparisons with dentist-moulded mouthpieces

To validate the quality of our mouthpiece in terms of both adhesion and impact on data quality, two participants received a dentist-moulded custom mouthpiece. Both participants underwent task-free fMRI scanning with no deliberate subject motion under a 2*2 design (with our mouthpiece or with dentist-moulded mouthpiece, with PMC applied or without PMC applied). Each permutation of conditions was scanned for 2 minutes with the same parameters as the 3mm EPI scans reported in the main experiment here.

We evaluated the effect of dentist-vs custom-moulded mouthpieces in three ways. Firstly, the tracking data from the camera were extracted and compared. This would allow for the quantification of the different amounts of involuntary motion induced by each mouthpiece respectively. Secondly, data from the scans where PMC was applied were processed using SPM. The magnitude of the realignment parameters from these scans is representative of the amount of motion that the PMC system has failed to correct for. Lastly, tSNR was calculated to test for any significant differences in fMRI data quality.

## RESULTS

### Participant comfort

Based on participant feedback, most participants felt that the mouthpiece was relatively comfortable, rating it an average of 3.1 (min: 0, max: 7) on a scale of 0 to 10 with 10 being extremely uncomfortable. 50% of the participants reported slight trouble swallowing. 94% of the participants indicated that they were willing to wear the mouthpiece for future scans, of which 84% expressed no reservations and 13% of the participants would only do so if it improved data quality. Ratings by participants were similar over the two sessions with mouthpieces, indicating that repeated use did not substantially alter the experience of the participants.

### Analysis of SPM motion parameters

The average integrated motion metric was obtained for each participant and plotted in Figure 3. All participants, except S07 and S11, demonstrate qualitatively similar motion profiles, with most residual motion for condition P-M+ and least residual motion for condition P+M+. There was a significant increase in motion between condition P-M+ (mean: 2.84mm/s) and condition P-M-(mean: 2.07mm/s) which provides evidence that the mouthpiece causes a slight increase in participant motion (p = 0.02, FFX permutation test). However, once PMC was applied in condition P+M+, the average motion metric showed a significant decrease (mean: 0.90mm/s) and was significantly lower than both condition P-M+ (p = 0.0001, FFX permutation test) and condition P-M-(p=0.0002, FFX permutation test). This indicates an overall beneficial impact of the PMC system on uncorrected head motion relative to a normal scan.

**FIGURE 3:** Plots of integrated motion metric of residual motion picked up by SPM post-processing for all participants. Most participants exhibit the common trend of least residual motion in Condition P+M+, followed by Condition P-M- and most residual motion in Condition P-M+. Error bars in the average integrated motion metric indicate standard error over participants.

### tSNR analysis of resting-state fMRI

The tSNR results were obtained from the resting-state fMRI (rsfMRI) run and are summarized in Figure 4. The histograms shown are pooled over all 15 participants and images in the top right hand corner of each plot show a representative slice through the tSNR map of a typical participant. There was a clear shift in distribution towards higher tSNR values in condition P+M+ (median tSNR: 73) compared to the other conditions (median tSNR: 65 and 66 for conditions P-M+ and P-M-respectively). A slight increase in voxels with low tSNR in the range of 10-40 was noticeable in condition P-M+ relative to condition P-M-, which may be a consequence of slight additional head motion in the presence of the mouthpiece. A paired permutation test was run using the median tSNR of the participants and no significant difference was observed between conditions P-M+ and P-M-(p=0.309, FFX permutation test) while condition P+M+ had significantly higher tSNR values relative to both condition P-M+ (p=0.043, FFX permutation test) and condition P-M-(p=0.022, FFX permutation test).

**FIGURE 4:**
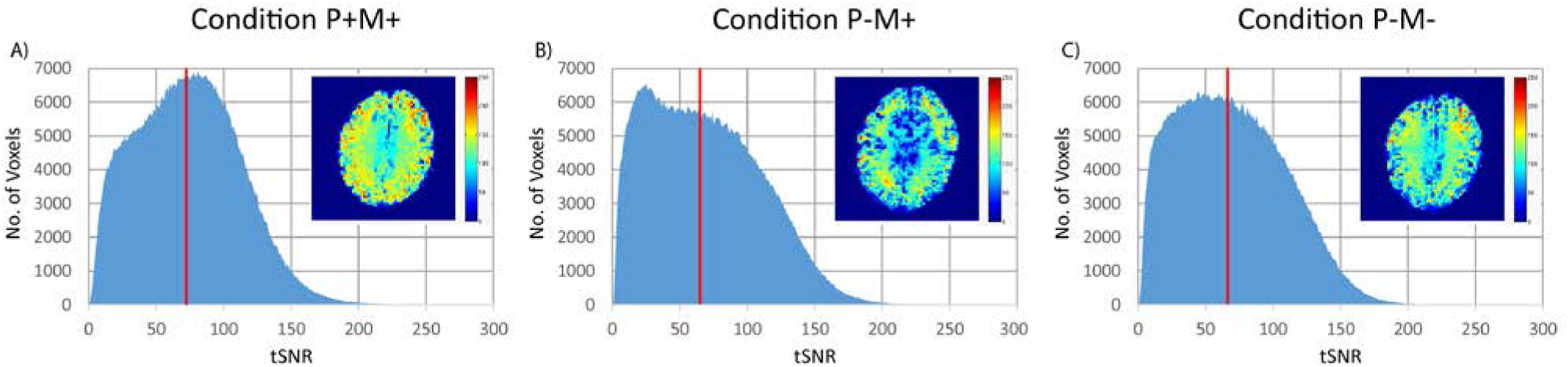
tSNR histograms for resting state fMRI comparing the three conditions. The tSNR values were pooled from all 15 participants into one histogram for each condition. The vertical red line indicates the median tSNR across all participants. A representative slice through the tSNR map of one participant is shown as an inset for each condition.

### Univariate Analysis using fCNR

Univariate analysis using the fCNR of voxels in V1 on the data from the main experiment showed comparable values for the individual conditions, at both 3mm (mean fCNR of conditions P+M+, P-M+ and P-M-: 1.1, 1.0 and 1.1 respectively) and 1.5mm resolution (mean fCNR of conditions P+M+, P-M+ and P-M-: 0.82, 0.81 and 0.84 respectively). The fCNR plots are shown in Supplementary Figure S5. Pairwise permutation testing showed no significant differences between the three conditions (all p>0.15, FFX permutation test).

### LDC Analysis

Results from the LDC analysis, plotted in Figure 5, demonstrated the benefits of PMC that depended on acquisition resolution. A three-way repeated measures ANOVA (Table 1) showed a main effect of region (F(2,14=10.559, p=0.0004). We interrogated this effect with post-hoc pairwise T tests after pooling data across PMC conditions and resolutions (Supplementary Table S6). This analysis confirmed that the radial bias ROI had a higher LDC than the entire V1 and the no radial bias ROI (p=0.0055 and p=0.0296, Tukey’s HSD test). Main effects of resolution and condition were non-significant (p>0.08).

**TABLE 1:** Repeated measures ANOVA results for LDC distance. * indicates p<0.05 (corrected for multiple comparisons, Tukey’s HSD test)

|  | Sum of Squares | DF | Mean Square Error | F | p |
| --- | --- | --- | --- | --- | --- |
| (Intercept) | 30.772 | 1 | 30.772 | 95.4790 | 1.24E-07 |
| Error | 4.5122 | 14 | 0.3223 |  |  |
| (Intercept):Resolution | 0.0043 | 1 | 0.0043 | 0.1275 | 0.7264 |
| Error(Resolution) | 0.4712 | 14 | 0.0337 |  |  |
| (Intercept):Condition | 0.2501 | 2 | 0.1251 | 2.7434 | 0.0817 |
| Error(Condition) | 1.2764 | 28 | 0.0456 |  |  |
| (Intercept):Region | 0.5208 | 2 | 0.2604 | 10.5590 | 0.0004* |
| Error(Region) | 0.6906 | 28 | 0.0247 |  |  |
| (Intercept):Resolution:Condition | 0.1977 | 2 | 0.0989 | 5.6633 | 0.0086* |
| Error(Resolution:Condition) | 0.4888 | 28 | 0.0175 |  |  |
| (Intercept):Resolution:Region | 0.0327 | 2 | 0.0163 | 2.9710 | 0.0676 |
| Error(Resolution:Region) | 0.1539 | 28 | 0.0055 |  |  |
| (Intercept):Condition:Region | 0.0177 | 4 | 0.0044 | 0.8313 | 0.5110 |
| Error(Condition:Region) | 0.2978 | 56 | 0.0053 |  |  |
| (Intercept):Resolution:Condition:Region | 0.0150 | 4 | 0.0038 | 1.2524 | 0.2996 |
| Error(Resolution:Condition:Region) | 0.1679 | 56 | 0.0030 |  |  |

**FIGURE 5:** Plot of normalized LDC distance per condition, resolution and ROI. The distance measures were averaged across all 15 participants. Error bars indicate standard error over participants. * indicates p<0.05 (corrected for multiple comparisons, Tukey’s HSD test).

The non-significant main effects of resolution and condition were moderated by a significant two-way interaction (F(2,14)=5.6633, p=0.0086), which suggests that the motion correction effects depended on data resolution. We interrogated this interaction further with post-hoc Tukey’s HSD tests after pooling data across regions. A significant improvement in LDC was observed for condition P+M+ relative conditions P-M+ at 1.5mm resolution (p=0.0086 respectively, Tukey’s HSD test, Supplementary Table S6). No significant differences were observed at 3mm resolution (all p>0.5, Tukey’s HSD test). No other two-way or three-way interactions were statistically significant (all p>0.06). Comparing conditions within each region and resolution showed that condition P+M+ had a significant improvement in LDC relative to condition P-M+ for all regions at 1.5mm resolution (p=0.022, p=0.045 and p=0.010 for the entire V1, regions with and without radial bias respectively, Tukey’s HSD test). Condition P+M+ also produced a significantly higher LDC relative to condition P-M- at 1.5mm resolution and looking at regions with no radial bias (p=0.031, Tukey’s HSD test).

These analyses indicate that PMC improved LDC effects, but this advantage was specific to high-resolution data. However, we were unable to show that this effect also depended on V1 sub-region.

### Classification Accuracy Analysis

fMRI data from the main experiment were used to train and test a SVM classifier to distinguish between the two orientations of the stimuli. Changes in the classification accuracy across conditions would be indicative of the effects of PMC on data quality. A summary plot is shown in figure 6.

**FIGURE 6:** Plot of the SVM classification accuracy results per condition, resolution and ROI. The accuracies were averaged across all 15 participants. Error bars indicate standard error over participants. * indicates p<0.05 (corrected for multiple comparisons, Tukey’s HSD test).

At 3mm resolution in the full V1 ROI, condition P+M+ had a classification accuracy of 89±2% (mean ±1 standard error) while conditions P-M+ and P-M-had a classification accuracy of 85±4% and 86±3% respectively. At a higher resolution of 1.5mm in V1, the mean classification accuracy dropped to 68±3%, 62±3% and 66±3% for conditions P+M+, P-M+ and P-M-respectively.

Using V1 sub-regions with expected radial bias yielded comparable classification accuracies of 90±2%, 88±3% and 82±3% at 3mm and 73±2%, 68±3% and 71±3% at 1.5mm for conditions P+M+, P-M+ and P-M-respectively. Performance in V1 sub-regions with no expected radial bias were overall lower than in the expected radial bias case (91±2%, 83±4% and 85±3% at 3mm and 70±2%, 66±3% and 68±2% at 1.5mm for conditions P+M+, P-M+ and P-M-respectively), with the exception of condition P+M+ at 3mm. This would be consistent with the notion that the no-radial-bias regions probed a finer scale spatial response, which would be more susceptible to motion effects. However, a repeated measures ANOVA, shown in Supplementary Table S7, showed only a significant effect of resolution (F(1,14)=135.3, p=1.39*10^-8^), with no significant main effects of condition and region, and no significant two-way or three-way interactions. Thus, despite a numeric benefit for PMC in all cases, analysis of SVM classification performance did not reveal a reliable advantage for PMC compared to the two control conditions.

### Simulation Results

We simulated how PMC modulates physiological noise levels, and assessed the sensitivity of each method to differences in fCNR by characterising the proportion of 100 simulated datasets that yielded a significant paired t-test (p<0.05) for a given fCNR pairing (Supplementary Figure 4). In this context, higher rejection probabilities for a given method indicates higher sensitivity to changes in fCNR. The simulation results demonstrate that LDC (Supplementary Figure 4, Panel A) is more sensitive to small changes in fCNR as compared to SVM (Supplementary Figure 4, Panel B) as seen from the higher rejection probability of the null hypothesis for the same change in fCNR. This is true for values of fCNR, thermal and physiological noise that are reflective of our real data, and this is consistent with our above results where the LDC is able to detect a significant improvement due to PMC, but not SVM. There is also a huge drop in sensitivity to changing fCNR at high fCNR for SVM, as seen by the fanning out of the region with high rejection probabilities at high fCNR of the SVM heatmap. This is due to ceiling effects since the classifier is classifying accurately for all iterations at high fCNR. This is similar to what we saw in our real data, with near ceiling performances at 3mm. Thus, this simulation demonstrates that higher sensitivity to PMC effects is expected for LDC relative to SVM.

### Comparison with dentist-moulded mouthpieces

In addition to the main experiment, we acquired dentist-moulded mouthpieces for two participants and compared the quality of the data acquired when using either mouthpieces. Initial comparisons for conditions where participants were told to remain as still as possible showed a slight increase in our integrated motion metric, measured via the mouthpiece, when participants were using our custom-moulded mouthpiece (0.58±0.30 mm/s, mean ±1 standard error) compared to the dentist-moulded mouthpiece (0.45±0.12 mm/s). However, this did not translate to an increase in residual motion as reflected in the realignment parameters after PMC had been applied: Our custom-moulded mouthpiece had an average residual integrated motion of 0.22±0.08 mm/s, compared to 0.24±0.01 mm/s for the dentist-moulded mouthpiece. The tSNR comparisons, as shown in Figure 7, also show no appreciable differences between the two mouthpieces.

**FIGURE 7:**
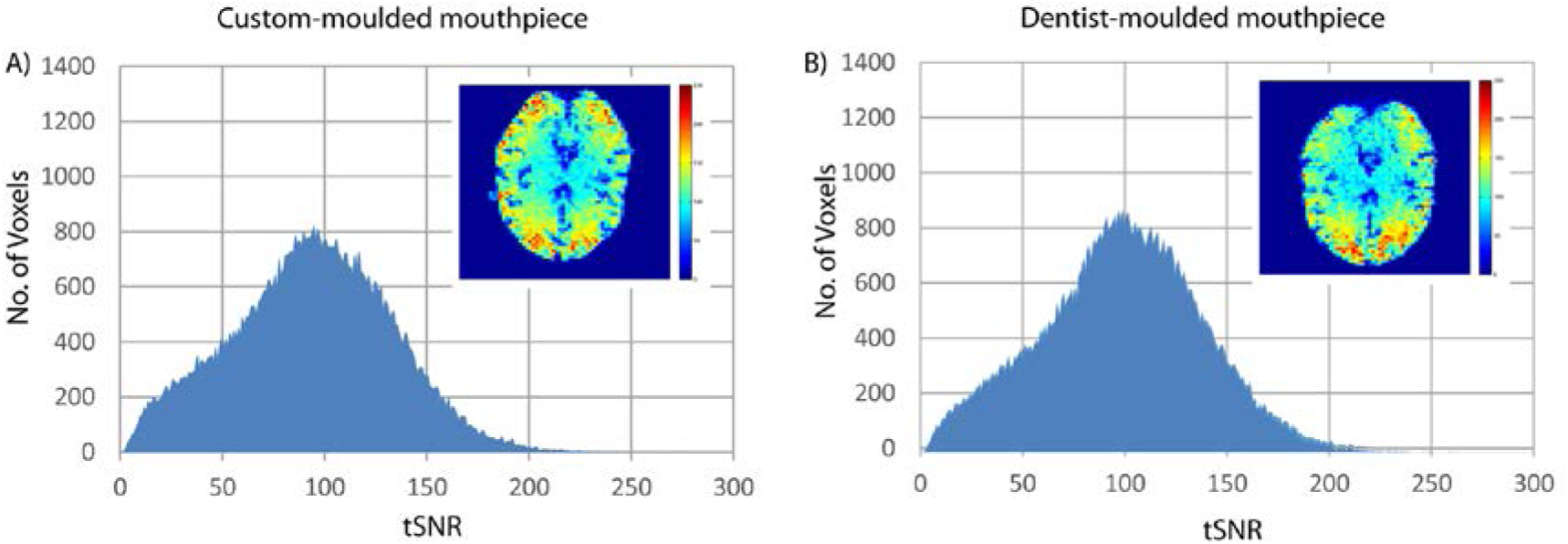
tSNR histograms for resting state fMRI comparing our mouthpiece against a dentist-moulded mouthpiece. The tSNR values were pooled from both participants into one histogram for each condition. A representative slice through the tSNR map of one participant is shown as an inset for each condition.

## DISCUSSION

Subject motion during an fMRI scan can significantly degrade image quality. Across two fMRI resolutions, we found evidence that PMC application improves data quality at higher resolutions (1.5mm). Given the increasing interest in laminar structures and fine scale functional specialization that are only visible at sub-millimetre resolution (Gilbert et al., 2010; Kok et al., 2016; Xing et al., 2012; Yacoub et al., 2008), it is likely that PMC will become crucial for advancement in these endeavours.

The tSNR data showed an improvement for PMC relative to the two control conditions, replicating previous studies (Todd et al., 2015). Univariate analysis using fCNR however showed no significant differences between conditions, unlike previous studies. This behaviour is expected because the univariate ROI analysis is only sensitive to average activation over a number of voxels within an ROI. Thus, the data would not be significantly affected by small motion artefacts. Since participants were instructed to keep as still as possible, it is likely to result in smaller differences across conditions compared to previous work, which used deliberate subject motion.

Analysis using the LDC revealed a benefit of PMC that appeared specific to high-resolution data. Post hoc multiple comparisons also showed that regions with radial bias generates the largest LDC, in line with studies by Freeman et al., 2013; Tong et al., 2010 which suggested that decoding in the visual cortex is strongly driven by radial bias rather than more fine-grained response patterns. There was also a numerical trend that improvements due to PMC were stronger in non-radial-bias V1 sub-regions, where the spatial activation patterns may have been expected to be more fine grained. However, this difference was not statistically significant when tested for using the three-way repeated measures ANOVA. In conclusion, we have demonstrated that the advantage of PMC is more apparent at higher resolutions, but we were unable to demonstrate a dependence on the expected spatial frequency of the activation patterns.

Analysis using SVM showed similar trends as the LDC results, with Condition P+M+ showing a consistent numerical improvement in classification accuracy over the other two conditions for both resolutions and all ROIs. However, these differences were not significant when tested for using ANOVA. We believe that this is a result of the lower sensitivity of SVM. Due to the limitations of SVM – namely discretization of results, rigid decision boundary across sub-runs and ceiling effects (see Introduction) – the SVM has more inherent variability and lower sensitivity and reproducibility across conditions (Walther et al., 2016). Moreover, due to the need to generate sufficient training data, each presentation block was modelled using an individual regressor. This increases the variance in the estimates for the decision boundary of SVM and contrast for LDC(see Supplementary Figure S2, Supplementary Table S3). We would expect these factors to translate into lower sensitivity of the SVM analysis to main effects and interactions present in the ANOVA. Thus, it is not unexpected to be able to detect the improvements of PMC using LDC, but not with SVM.

Our simulation results also support our findings reported above. We showed that in the regime of our data, LDC is more sensitive to fluctuations in fCNR, and thus more likely to detect improvements by PMC compared to SVM.

Our results showed that PMC is particularly important for studies at higher resolution and studies that require accurate voxel registration. For robust activations and simple analysis, such as fCNR of the primary visual cortex, there would appear to be little discernible benefit of PMC, in line with the results obtained by Zaitsev et al., 2016. For high-resolution multivariate pattern decoding analyses where accurate voxel registration across time is essential, there was a clear benefit of PMC. There is also the question of whether a dentist-moulded mouthpiece is more stable and comfortable than our custom-built mouthpiece. We found, at least on two participants, that a dentist-moulded mouthpiece yielded similar data quality to our custom-moulded mouthpiece. While both participants felt slightly more comfortable with the dentist-moulded mouthpiece, this was not reflected in the post-processing motion parameters nor in tSNR estimates. This suggests that instead of motion arising due to the discomfort, the cause of motion is the inherent presence of an attachment in the mouth, independent of type of attachment used. This also demonstrated that our commercially available solution to marker attachment shows comparable performance to the more expensive mouthpiece utilized by other sites.

The mouthpiece utilized in this experiment has the added benefit of accessibility and convenience, since it is relatively inexpensive and can be moulded on the spot a few minutes prior to the actual experiment. Moreover, the presence of a dentist is not required. Based on participant feedback, most people did not find the mouthpiece uncomfortable and an overwhelming majority indicated that they were willing to use the mouthpiece for the purpose of data quality improvement.

Our study supports and extends previous studies on PMC for fMRI. Rotenberg et al., 2013 and Schulz et al., 2014 both demonstrated improvements in data quality when subject motion, both deliberate and task-correlated, was present. Zaitsev et al., 2016 conducted an experiment with a similar paradigm but did not find any significant differences, likely due to the smaller sample size, choice of robust activation pattern and poor marker adhesion. As far as the authors are aware, the present study is the first showcasing significant benefits of applying PMC to task fMRI when participants have been instructed to remain as still as possible.

There are of course limitations to our study. Firstly, we have only employed 2D EPI sequences. However, other studies have shown similar results with 3D EPI (Todd et al., 2015) and diffusion weighted imaging (Herbst et al., 2012), albeit using different forms of marker attachment. Secondly, it is important to note that PMC implementation does not create the absolute gold standard for data quality because it is unable to correct for head motion through inhomogeneous B0 and B1 fields. B0 field distortions due to susceptibility changes at tissue boundaries can cause signal dropouts and geometric distortions (Hutton et al., 2013). This could be addressed using complimentary strategies together with PMC implementation (Glover et al., 2000; Lutti et al., 2013).

## CONCLUSION

With an increasing focus on higher fields and higher resolutions, subject motion during fMRI will remain a pertinent problem. Results acquired from this PMC system, both in this paper and elsewhere, have shown great promise for minimizing the negative impact of subject motion by constantly updating the scanner on the latest co-ordinates of the participant’s head position. Our results demonstrated that PMC is beneficial even when participants are instructed not to move, and that this benefit appears specific to high-resolution fMRI acquisitions. The custom mouthpiece utilized in our setup can greatly reduce the cost and inconvenience for marker attachment, potentially leading to more widespread adoption of the PMC system.

## ACKNOWLEDGEMENTS

The authors are grateful to all the radiographers at MRC Cognition and Brain Sciences Unit for their assistance in performing the scans and to the volunteers for participating in the experiment. R.N.H is supported by Medical Research Council Programme Grant (SUAG/010 RG91365).

## CONFLICT OF INTEREST

The authors have no conflict of interest to declare.

